# Cardio-audio synchronization elicits neural and cardiac surprise responses in human wakefulness and sleep

**DOI:** 10.1101/2022.03.03.482861

**Authors:** Andria Pelentritou, Christian Pfeiffer, Sophie Schwartz, Marzia De Lucia

## Abstract

The human brain can infer temporal regularities in auditory sequences with fixed sound-to-sound intervals and in pseudo-regular sequences where sound onsets are locked to cardiac inputs. Here, we investigated auditory and cardio-audio regularity encoding during sleep, when reduced vigilance may result in altered bodily and environmental stimulus processing. Using electroencephalography and electrocardiography in healthy volunteers (N=26) during wakefulness and sleep, we measured the response to unexpected sound omissions within three auditory regularity conditions: synchronous, where sound and heartbeat are temporally locked, isochronous, with fixed sound-to-sound intervals, and a control condition without specific regularity. During wakefulness and all sleep stages, the cardiac signal following sound omissions exhibited a deceleration over time in the synchronous condition only. At the neural level, both the synchronous and isochronous sequences gave rise to a neural omission response in wakefulness and N2 sleep. Regularity encoding in the synchronous and isochronous conditions was characterized by a modulation of the background oscillatory activity in N2 sleep, outlining a possible mechanism through which the brain aligns periods of high neuronal excitability to the expected sound onset. The violation of auditory and cardio-audio regularity elicits cardiac and neural surprise responses across vigilance stages.

**Significance Statement:** Across vigilance states, the human brain can generate predictions about the future based on past sensory regularities. While this is evident for environmental stimuli, the role of bodily signals in forming sensory prediction remains unknown. Here, we show that the human brain utilizes the temporal relationship between cardiac and auditory inputs in order to anticipate upcoming sounds during wakefulness and sleep. After presenting sounds in synchrony with the ongoing heartbeat, a sound omission elicited both a heartbeat deceleration and a prediction error signal as measured by the electroencephalographic response. Heartbeat signals support auditory regularity encoding during sleep and wakefulness, highlighting one mechanism for optimizing the detection of unexpected stimuli by taking advantage of the continuously monitored cardiac signals.

## Introduction

The processing of auditory regularity is a basic brain mechanism that enables the rapid detection of unexpected stimuli which can persist even in altered states of consciousness such as sleep and coma (1). The mechanism underlying auditory regularity encoding has mostly been studied by investigating the neural responses to deviant sounds interrupting a sequence of repeated standard stimuli as reported in healthy human wakefulness (2, 3), during sleep (*e.g.* (4–6)), and in disorder of consciousness patients (7–9). This rudimental component of auditory discrimination, known as mismatch negativity (MMN), has often been interpreted in the framework of the predictive coding theory (2, 3, 10, 11). According to this theory, the MMN may arise from the contribution of multiple, non-exclusive mechanisms including repetition suppression in response to frequent stimuli and generation of a ‘prediction error’ following an unexpected mismatch between the predicted and presented stimuli. The predictive nature of the neural responses to violations within regular auditory sequences has received experimental support from studies on unexpected omissions (12–16) wherein the top-down prediction is not confounded by the neural response to deviant sound stimuli.

In addition to external stimuli, the human brain receives internally generated bodily signals, which constitutes a continuous source of sensory inputs. Simultaneous environmental and bodily information may compete for neural resources and influence their respective processing (17). Accordingly, previous studies have shown that bodily signals and their associated neural representation modulate perception and cognition (18, 19). In particular, the neural processing of cardiac signals, as measured by heartbeat evoked potentials (HEPs; (20)), may determine whether a stimulus is consciously perceived (21), influence emotional processing (22–24) and could account for the first person perspective in perceptual experience (25). In addition, heartbeat processing can be used to measure interoceptive ability (*i.e.* the ability of sensing the inner body state), which in turn can influence affective control, physical and mental wellbeing in health and a variety of clinical conditions (26, 27).

In this context, it is also plausible that the rhythmic information from the heartbeat could modulate the processing of temporally-organized sequences of external sensory signals. Recent experimental paradigms have demonstrated that auditory sequences locked to the ongoing heartbeat generate an auditory temporal prediction even in the absence of fixed sound-to-sound intervals (28–30). Whereas such prediction across interoceptive and exteroceptive signals may arise during the awake state, it is unclear whether it may be observed during sleep when the processing of external information and its temporal structure is reduced compared to wakefulness (31). Previous studies in sleep have investigated interoceptive and exteroceptive stimulus processing independently. First, heartbeat stimulus processing across sleep stages (32–34) and its modulation in patients with sleep disorders in comparison to healthy controls (35, 36) suggest some preservation of interoceptive signal processing in sleep. Second, evidence of rudimental sensory stimulus processing in sleep is documented in a large body of literature (*e.g.* (6, 37–39)).

Here, we investigated whether the heartbeat and auditory signals are integrated and inform auditory regularity encoding. Addressing this question is important in light of previous evidence showing that the neural processing of cardiac signals can boost the integration of conscious percepts (20, 21), while the potentially beneficial effect of the heartbeat on external stimulus processing during sleep, when conscious awareness of the external environment is at its lowest, has never been shown. Specifically, we investigated whether detecting regularity in trains of auditory stimuli may benefit from their temporal alignment with the ongoing heartbeat in sleep and wakefulness. We hypothesized that, as the brain gradually disconnects from the environment during sleep, bodily signals may become increasingly critical for informing auditory regularity encoding and the detection of unexpected violation of such regularities. Healthy volunteers underwent two separate sessions of simultaneous electrocardiography (ECG) and electroencephalography (EEG) recordings during wakefulness and full night sleep.

Participants passively listened to two possible varieties of auditory regularities (Fig. 1). In the first condition, sounds were presented at a fixed short delay relative to the ongoing heartbeat (synchronous or synch condition), in the second, sounds were presented at a fixed sound-to-sound interval (isochronous or isoch condition) and a third condition, where sounds were presented without any specific regularity (asynchronous or asynch condition) served as a control condition. To assess the preservation of regularity encoding in all conditions, we measured the cardiac responses (from ECG) and the neural responses (from EEG) to unexpected omissions interspersed within the auditory sequences. Upon sound omissions in wakefulness and sleep, we expected prediction error generation in both the synch and isoch conditions as a consequence of the cardio-audio regularity and auditory regularity encoding. As both the cardiac and the neural responses during sound omission may carry the signature of this prediction error generation (29, 40), we investigated the ECG and EEG responses to unexpected omission across different conditions of temporal alignments between cardiac and auditory signals.

**Fig. 1.**
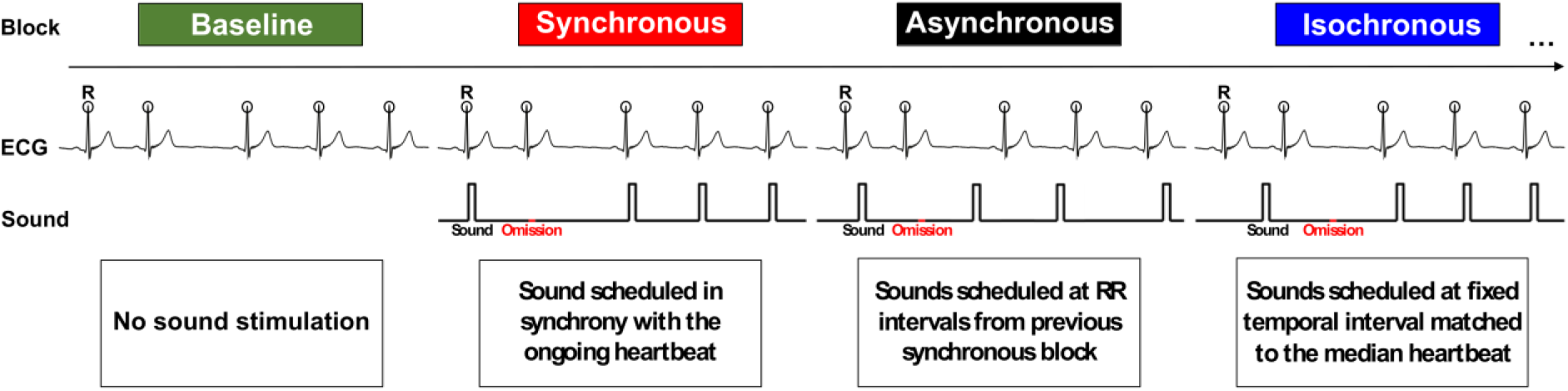
Experimental paradigm overview. The data acquisition was organized in four block types, administered in pseudo-random order. No auditory stimulation was performed during the Baseline condition. In the Synchronous condition, the R peaks (circles) were detected online in the ECG signal and the sounds were administered at a fixed 52 ms R-to-sound. In the Asynchronous condition, sound-to-sound intervals from a preceding Synchronous blocks were administered resulting in variable R-to-sound and sound-to-sound intervals. In the Isochronous condition, the median RR interval from a preceding Synchronous block was computed and served as the fixed sound-to-sound interval. Sound omissions occurred pseudo-randomly for 20% of trials within each block.

## Results

### 1. Participants and sleep characteristics

Twenty-six healthy volunteers participated in the study (14 female; 1 left-handed; mean age: 27 years, range: 20-35 years) and each took part in a wakefulness and sleep recording session. All 26 participants were included for wakefulness while the sleep dataset included 25 participants (13 female; 1 left-handed; mean age: 26 years, range: 20-35 years) due to malfunctioning equipment during the sleep session for 1 participant. The sleep characteristics recorded during the experimental night across the eligible population (N=25) are summarized in Table 1.

**Table 1.** Group averaged sleep characteristics. Total Bed Time calculated based on Lights Off and Lights On time points. Total Sleep Time computed as the period between the Sleep Onset and final awakening, uncovered by the sleep staging, excluding wakefulness periods but including possible micro-arousal periods during the night. Percentages of sleep stage periods (N1, N2, N3, REM) during the session were computed in relation to the Total Sleep Time. Sleep Efficiency (%) = Total Sleep Time/Total Bed Time for each participant×100. SD = Standard Deviation.

| N=25 | Mean | SD |
| --- | --- | --- |
| **Total Bed Time (min)** | 420.3 | 93.8 |
| **Total Sleep Time (min)** | 302.4 | 107.9 |
| **Sleep Onset (min)** | 24.6 | 19.1 |
| **Sleep Efficiency (%)** | 69.8 | 12.9 |
| **N1 (%)** | 12.9 | 4.5 |
| **N2 (%)** | 57.4 | 6.9 |
| **N3 (%)** | 15.5 | 8.8 |
| **REM (%)** | 14.1 | 5.5 |

### 2. Cardiac omission response

We investigated whether regularity violation upon omission of expected sounds could elicit a cardiac response across vigilance states. We analyzed heartbeat changes based on the RR intervals extracted from the ECG in response to sound omissions as a function of auditory conditions (synch, asynch, isoch). We included all 26 participants for the wakefulness session, 15 for N1 sleep, 24 for N2 sleep, 18 for N3 sleep, and 15 for REM sleep.

#### 2.1. Auditory condition contrast

We compared average RR intervals before, during and after omissions using two-way repeated measures ANOVAs with factors Auditory Condition (synch, asynch, isoch) and Trial Order (one trial before, trial during, first trial after, second trial after sound omissions) in the wakefulness and sleep sessions. To take full advantage of the available data in each vigilance state (Table S1-S2), separate repeated measures ANOVAs were performed for each vigilance state. These analyses were conducted on normalized RR intervals (by subject-wise division of each of the investigated average RR intervals by the average RR interval prior to sound omission), so that the results would not be attributable to inter-subject variability in RR intervals (Fig. 2).

**Fig. 2.**
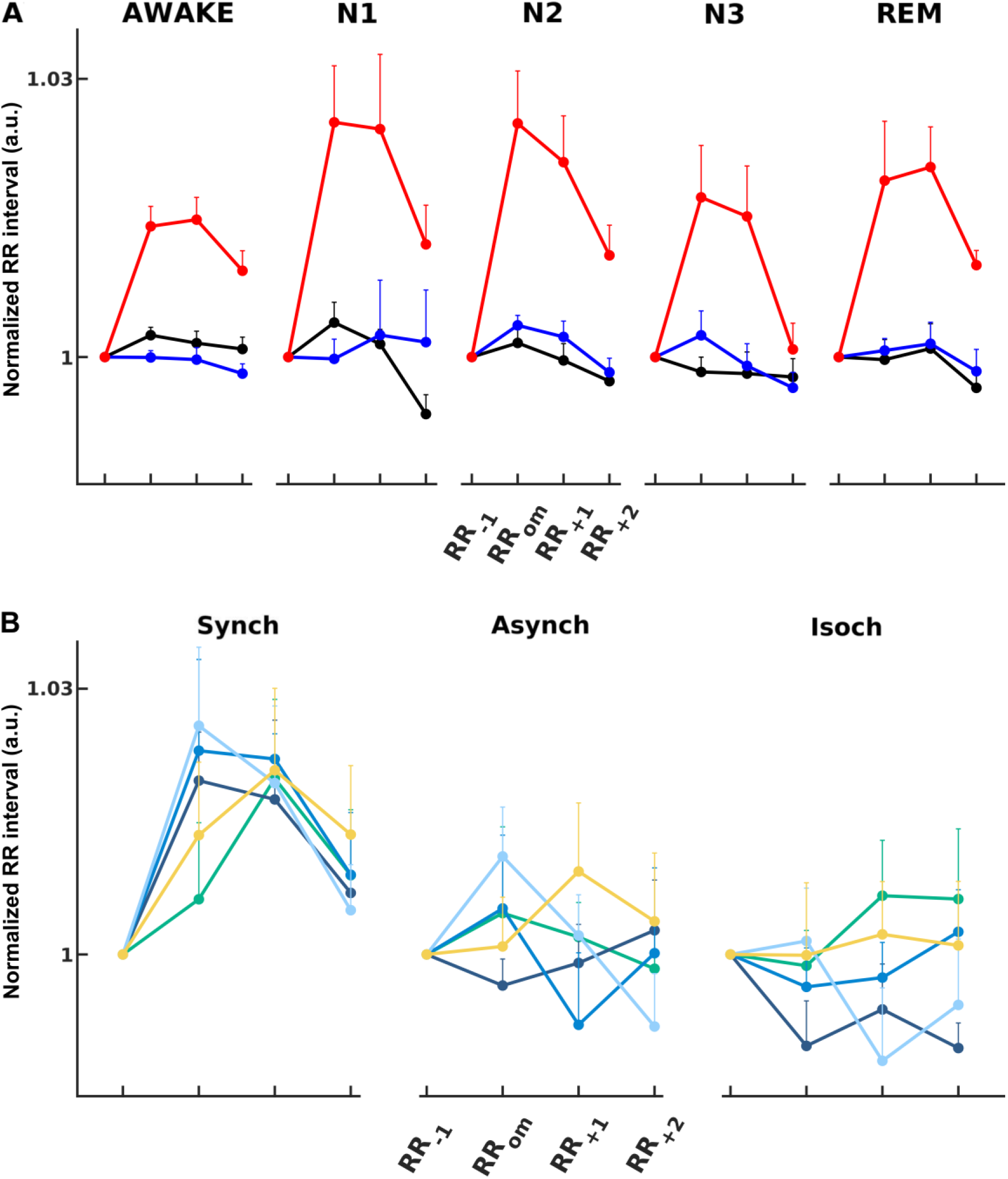
Cardiac omission response in wakefulness and sleep. (A) Grand averaged normalized RR intervals prior (RR_-1_), during (RR_om_), and following (RR_+1_ and RR_+2_) sound omissions across auditory conditions (synch: red line; asynch: black line; isoch: blue line) in the wakefulness session (AWAKE) and all sleep stages (N1, N2, N3, REM). (B) Grand averaged normalized RR intervals for participants with sufficient data across all vigilance states, prior (RR_-1_), during (RR_om_), and following (RR_+1_ and RR_+2_) sound omissions in the wakefulness session (AWAKE, yellow) and all sleep stages (N1, light blue; N2, darker blue; N3, darkest blue; REM, green) for each auditory condition. Error bars indicate the SEM (upper limit). Statistical analyses highlighted a heart rate deceleration (p<0.05) upon sound omission in the synch condition across all vigilance states.

Overall, in wakefulness and sleep, we observed that cardio-audio synchronization led to a heartbeat deceleration, particularly pronounced on the first heartbeat after sound omission (RR_om_) and persistent for at least two heartbeats (RR_+1_ and RR_+2_). This deceleration in the heart rate upon sound omission was absent or inconsistent in the asynch and isoch conditions.

Statistical analysis confirmed this difference across auditory conditions as follows. 3×4 repeated measures ANOVAs revealed a significant interaction of Auditory Condition x Trial Order for all vigilance states (Fig. 2A; AWAKE: F_(3,4)_=14.1, p=1.0×10^-12^; N1: F_(3,4)_=6.6, p=8.9×10^-6^; N2: F_(3,4)_=11.6, p=1.6×10^-10^; N3: F_(3,4)_=6.2, p=1.6×10^-5^; REM: F_(3,4)_=6.1, p=2.2×10^-5^) with a main effect of Auditory Condition (AWAKE: F_(3,1)_=27.2, p=10.0×10^-9^; N1: F_(3,1)_=11.3, p=2.5×10^-4^; N2: F_(3,1)_=14.4, p=1.4×10^-5^; N3: F_(3,1)_=9.0, p=8.9×10^-4^; REM: F_(3,1)_=17.6, p=1.1×10^-5^) and a main effect of Trial Order (AWAKE: F_(1,4)_=20.1, p=1.2×10^-9^; N1: F_(1,4)_=5.3, p=0.004; N2: F_(1,4)_=18.4, p=7.0×10^-9^; N3: F_(1,4)_=3.5, p=0.023; REM: F_(1,4)_=7.5, p=4.1×10^-4^). Permutation testing followed by Wilcoxon signed-rank tests evaluated the distribution of post-permutation F values against the original F values and confirmed the significance of the main effects and interactions (p<0.0005). Post-hoc paired Wilcoxon signed-rank tests with Bonferroni correction for multiple comparisons across conditions corroborated that in the synch condition (Fig. 2A, red line), omissions elicited a long-lasting heart rate deceleration, with higher RR interval during (AWAKE: p=1.2×10^-5^; N1: p=1.2×10^-5^; N2: p=2.4×10^-5^; N3: p=0.001; REM: p=8.5×10^-4^) and immediately after (AWAKE: p=1.5×10^-5^; N1: p=6.1×10^-5^; N2: p=2.1×10^-5^; N3: p=0.003; REM: p=6.1×10^-5^) the omission than before the omission across all vigilance states.

#### 2.2. Vigilance State Contrast

To investigate whether the heartbeat deceleration was modulated by vigilance state and auditory conditions (Fig. 2B), we computed a three-way repeated measures ANOVA with factors Vigilance State (AWAKE, N1, N2, N3, REM), Auditory Condition (synch, asynch, isoch), and Trial Order (one trial before, trial during, first trial after, second trial after sound omission), including only participants who had sufficient data in all vigilance states (N=6; Table S2).

The ANOVA revealed a significant main effect of Auditory Condition (F_(1,3,1)_=27.5, p=1.1×10^-11^), due to overall higher RR interval values (*i.e.* enhanced deceleration) during the synch condition, and a main effect of Trial Order (F_(1,1,4)_=3.9, p=0.009) but no main effect of Vigilance State (p>0.05). Critically, replicating the analyses on each vigilance state separately (Fig. 2A), the interaction effect of Auditory Condition and Trial Order was also significant (F_(1,3,4)_=4.2, p=4.0×10^-4^). Permutation testing followed by Wilcoxon signed-rank tests verified the significance of the main effects and interaction (p<0.0005). The interactions Vigilance State x Auditory Regularity, Vigilance State x Trial Order, and the triple interaction across all factors (Vigilance State x Auditory Regularity x Trial Order) were not significant (p>0.05). This pattern of results suggests that the heart rate deceleration upon sound omission in the synch condition occurred during wakefulness and sleep and to a similar extent across vigilance states.

### 3. Neural omission response

In the EEG analysis, 23 participants were eligible for wakefulness (AWAKE) and N2 sleep as well as 12 for N1 sleep, 14 for N3 sleep, and 13 for REM sleep (Table S1-S2). Since the sample size estimation for obtaining statistically significant results in the EEG comparisons (see Methods. Sample size estimation) revealed a minimum sample size of 17 participants, we did not compare the EEG evoked responses for N1, N3 or REM sleep wherein the sample size criterion was not met.

#### 3.1. Auditory evoked response

Sounds elicited auditory evoked potentials in all sleep stages (AEPs, Fig. 3) with no differences in the AEPs between the isoch and asynch conditions (cluster permutation statistical analysis, p>0.05; two-tailed) in wakefulness and N2 sleep. We refrained from performing these comparisons in the synch condition as the AEPs in that case were contaminated with the response to the heartbeat.

**Fig. 3.**
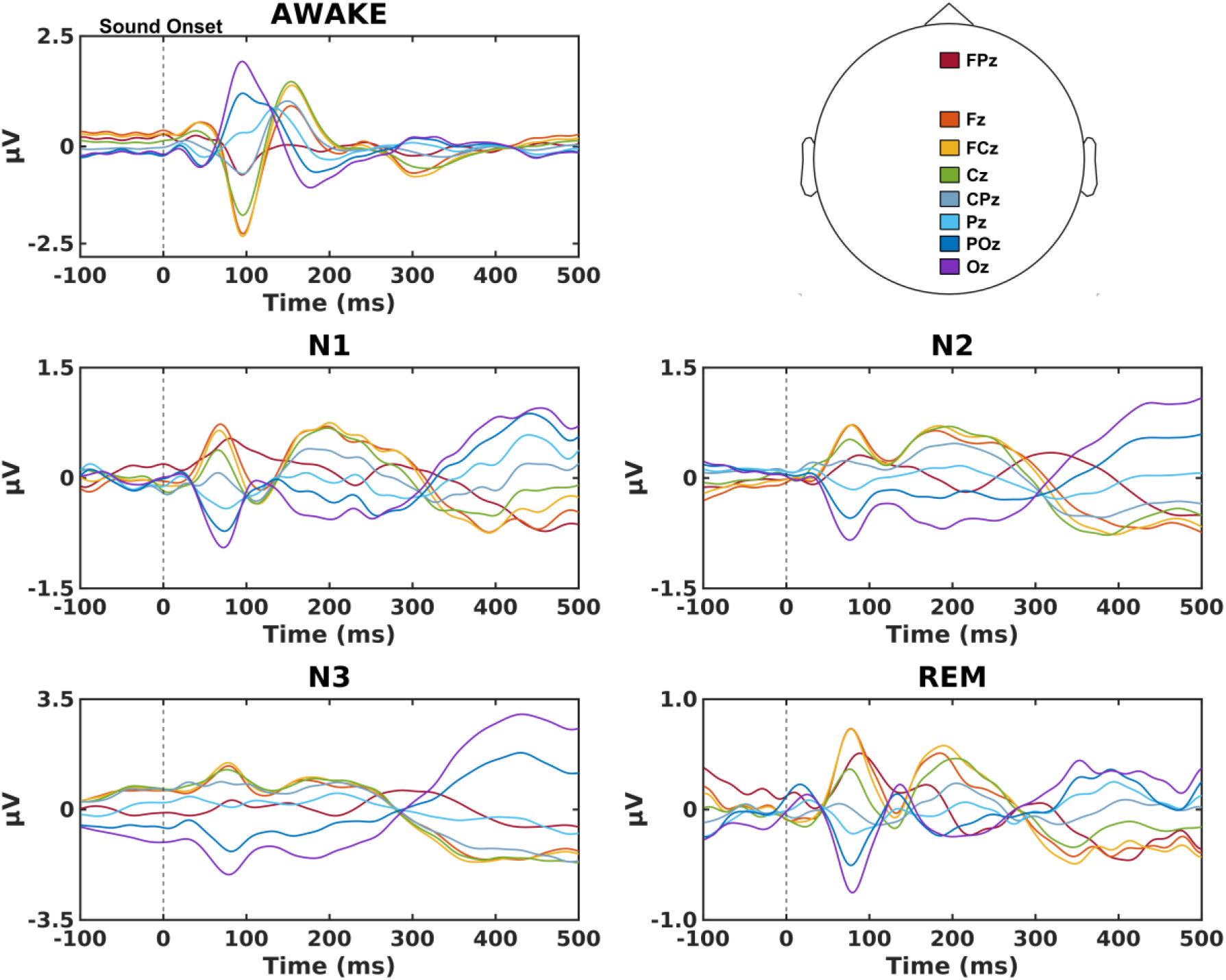
AEPs in wakefulness and sleep. Grand averaged AEPs in the isoch condition on EEG electrodes located along the midline in wakefulness (AWAKE) and all stages of sleep (N1, N2, N3, REM). AEPs in wakefulness display the expected electrophysiological signatures such as the N100 as the first and highest in amplitude component and the expected shift to later P200 and N550 components across sleep stages. Asynch condition AEPs are not depicted due to the high similarity to the isoch condition (confirmed by cluster permutation statistical analysis, p<0.05, two-tailed, showing no statistically significant differences in all vigilance states) and synch condition AEPs are not depicted as they would be superimposed with the heartbeat evoked potential.

#### 3.2. Omission response in cardio-audio regularities

To investigate the effect of cardio-audio synchronicity on omission responses, we derived heartbeat-evoked potentials during sound omissions (OHEPs) in the synch and asynch conditions for wakefulness and N2 sleep. Average OHEPs were calculated by extracting epochs from the continuous EEG recordings that were time-locked to the first R peak of the ECG signal during omissions. As an additional control condition for this analysis, we extracted a random selection of R peaks in the ECG signal to derive average HEPs from the continuous EEG recordings in the baseline condition.

In the wakefulness session, we expected to observe differences in the HEPs when comparing the synch to asynch and the synch to baseline conditions, as a consequence of the predictability of sound onset in the synch condition, based on the fixed delay between R peaks and sounds (29). For the synch *vs* asynch comparison (Fig. 4A), the cluster-based permutation test (p<0.05, two-tailed) revealed a significant negative cluster (p=0.033, Cohen’s d=1.055) at 165 ms to 218 ms following R peak on anterior-central scalp electrodes. Similar results were observed in the synch *vs* baseline comparison (Fig. S1A) while the asynch *vs* baseline comparison showed no significant differences (Fig. S1B).

**Fig. 4.**
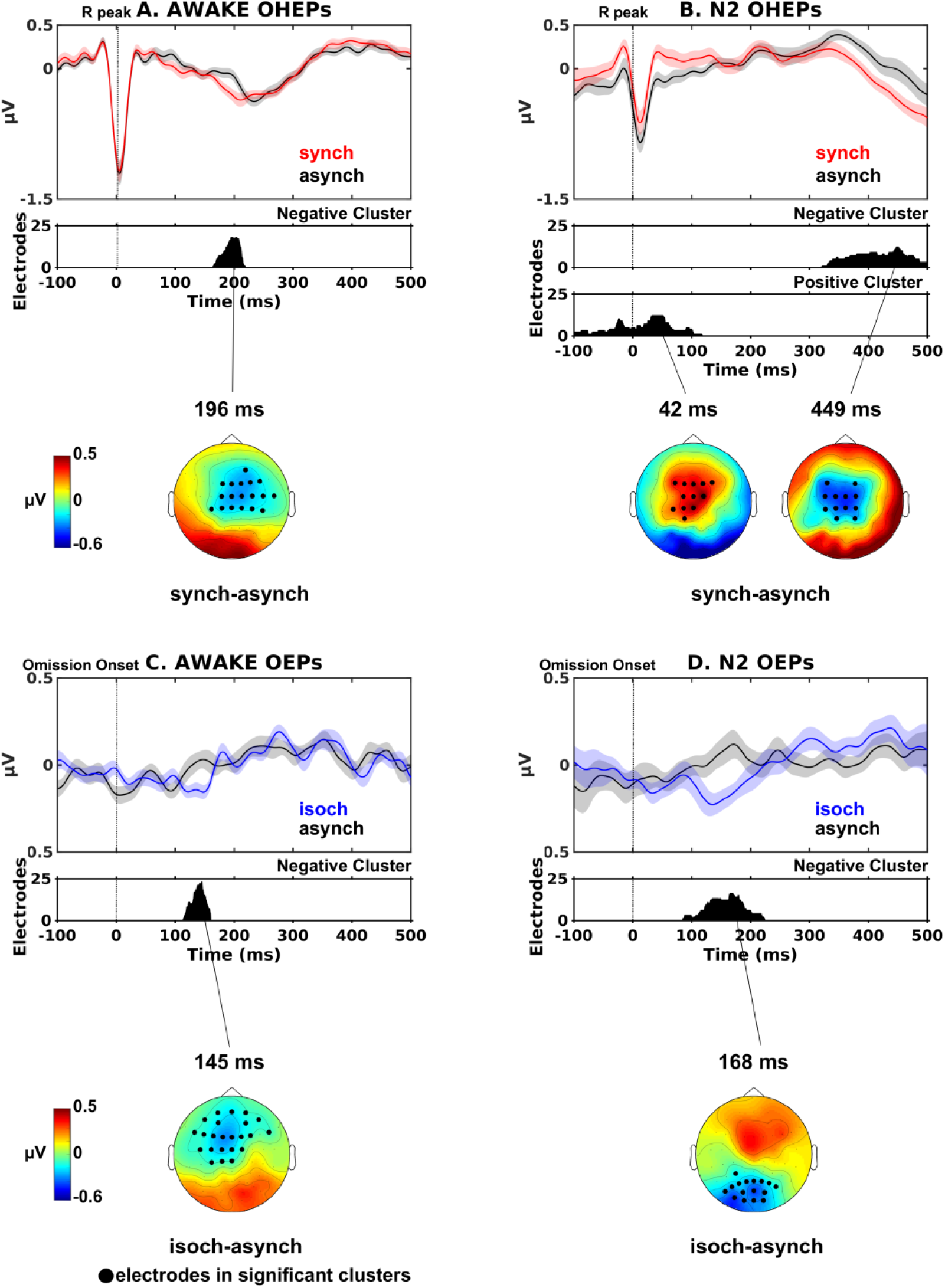
Neural omission response in wakefulness and N2 sleep. OHEPs (time-locked to the R peak) and OEPs (time-locked to mean SS interval) comparisons during sound omissions for the isoch (blue lines) and synch (red lines) conditions compared to the asynch condition (black lines) during wakefulness (AWAKE) and N2 sleep (N2). Top panels show grand averaged (N=23) EEG waveforms averaged over all electrodes in the significant negative clusters. Shaded regions indicate ±SEM across participants. Middle panels display the number of electrodes within each significant cluster derived from cluster permutation statistical analysis (p<0.05, two-tailed). Bottom panels demonstrate topography differences at cluster peaks with significant electrodes highlighted in black. Significant differences are observed between the synch and asynch OHEPs in wakefulness (A) and N2 sleep (B), suggesting that the synch cardio-audio regularity induces a modulation of OHEPs both in wakefulness and N2 sleep. Significant differences between the isoch and asynch condition OEPs in wakefulness (C) and N2 sleep (D) demonstrate that the isoch regularity produces an expectation of incoming sounds observed as a MMN response at ∼150 ms, in wakefulness and N2 sleep.

In N2 sleep, we found differences between the OHEPs of the synch and asynch conditions (Fig. 4B) at –99 ms to 117 ms (p=0.049, Cohen’s d=0.836) and at 322 ms to 500 ms (p=0.017, Cohen’s d=0.622) following R peak onset; both clusters were localized to central electrodes. In N2 sleep, we did not find significant results neither in the comparison of synch *vs* baseline nor in the contrast of asynch *vs* baseline.

As sound omissions are unexpected, the early significant difference between the synch and asynch conditions observed only during N2 sleep could be explained by changes in the background oscillatory activity during this vigilance state (addressed below in section 4. Slow oscillation analysis). This was further confirmed when contrasting the synch AEPs *vs* OHEPs in N2 sleep, a contrast that yielded one positive cluster (p=4.0×10^-4^) between 208 ms and 500 ms on central electrodes and one negative cluster (p=4.0×10^-4^) between 203 ms and 500 ms on peripheral electrodes, suggesting that this early differential response in the synch *vs* asynch comparison could relate to auditory stimulus anticipation in response to a heartbeat.

Overall, our results indicate that similar to a fixed sound-to-sound interval, the fixed heartbeat-to-sound interval induced auditory prediction observed as a neural surprise MMN-like response upon regularity violation during omissions in both wakefulness and N2 sleep.

#### 3.3. Omission response in auditory regularities

As a further validation of the existence of an auditory predictive processing mechanism during wakefulness and sleep, we tested whether fixed sound-to-sound intervals induced an expectation of upcoming auditory stimuli, violated upon omission. To do so, we derived omission responses during sound omissions in the isoch and asynch conditions in wakefulness and N2 sleep. Average sound-based omission evoked potentials (OEPs) were calculated by extracting epochs from the continuous EEG recordings that were time-locked to the average sound interstimulus interval. A random selection of epochs were extracted from continuous baseline recordings, such that the latencies between epoch onset and closest heartbeat (*i.e.* R peak) were matched to the trial onsets in the sound-based isoch and asynch conditions at the single-trial level.

Based on previous reports of neural responses to omissions within regular auditory sequences during wakefulness (*e.g.* (12)), here, we expected central negativity in the isoch condition (where temporal regularity existed in sound stimuli) between 100 ms and 250 ms. In the wakefulness session, the cluster-based permutation test (p<0.05, two-tailed) comparing the OEPs in the isoch *vs* asynch condition yielded a significant negative cluster (p=0.033, Cohen’s d=0.919) at 114 ms to 159 ms following expected sound onset in anterior-central scalp electrodes (Fig. 4C). The existence of an omission response in the isoch condition was further confirmed by the isoch *vs* baseline comparison (Fig. S1C). Despite the absence of regularity in the asynch condition, the asynch *vs* baseline comparison in wakefulness revealed a significant negative cluster approximately at the expected sound onset (∼0 ms; Fig. S1D). This last result indicates some degree of prediction of upcoming sounds in the asynch sequence during wakefulness despite the absence of temporal expectation, plausibly due to the pseudo-regularity of the auditory sequence.

In N2 sleep, the isoch *vs* asynch OEP comparison revealed significant differences that were largely similar to wakefulness, at least in terms of latency relative to expected sound onset. In more detail, statistical evaluation of the isoch and asynch condition differences (Fig. 4D) identified a significant negative cluster (p=0.043, Cohen’s d=0.660) at 85 ms to 223 ms following expected sound onset localized to posterior-central scalp electrodes. Different to wakefulness, no significant clusters were identified when contrasting the isoch or asynch to the baseline condition in N2 sleep.

Our results in wakefulness and N2 sleep suggest that in the isoch condition, the fixed sound-to-sound interval induced an expectation of upcoming sounds and resulted in a neural surprise response upon violation of the regularity rule during omissions.

### 4. Slow oscillations analysis

As outlined above (Fig. 4B), the analysis of OHEPs during N2 sleep (N=23) revealed that cardio-audio synchronization (compared to the asynch condition) gave rise to a central early onset positivity (at –99 ms to 117 ms) and late onset negativity (at 322 ms to 500 ms). This effect is reminiscent of the up and down states of slow oscillations (SOs; 0.5-1.2 Hz oscillations) during N2 sleep (41, 42). This interpretation was confirmed by the analysis of the band-passed EEG data at 0.5 Hz to 4 Hz (delta band; reflecting the range of slow wave frequencies) which produced a highly similar profile for the synch *vs* asynch OHEPs during N2 sleep with an early onset central positivity difference (p=0.030) between –99 ms and 125 ms and a late onset central negativity difference (p=0.023) between 259 ms to 500 ms following heartbeat onset.

In light of this observation and since sound presentations are known to alter the background oscillatory activity in sleep, notably the SOs during NREM sleep (39, 43, 44), we investigated whether the three auditory conditions had differential effects on the ongoing oscillations during N2 sleep. We selected the SO positive half-wave peak latency at electrode Cz (Fig. 5AB) as the representative SO latency and computed the median sound-to-SO latency for all auditory conditions and median R peak-to-SO latency for all auditory conditions and the baseline.

Statistical assessment using an 1×3 repeated measures Friedman test on the mean sound-to-SO latencies yielded significant differences across auditory conditions (Fig. 5C; χ^2^ =7.9, p=0.019). Post-hoc Wilcoxon signed-rank tests confirmed lower latencies in synch compared to asynch (p=0.009) and in isoch compared to asynch (p=0.026). These results suggested a possible readjustment of the SOs with respect to sound onset depending on the regularity condition (Fig. 5C) since when regularity was present, either in the synch or isoch condition, SOs tended to align to the sound onset. Because of the fixed RS delay in the synch condition, the alignment between sound onset and SOs was also reflected in a lower R peak to SO peak in the synch compared to the asynch and isoch conditions (Fig. 5D). In order to rule out that this SO readjustment was due to a specific relation between R peak and SO irrespective of sound presentation, we carried out a 1×4 repeated measures Friedman test on the mean R peak-to-SO latencies which revealed significant differences across conditions (Fig. 5D; χ^2^ _(1,4)_=7.4, p=0.062), suggesting that potential readjustment of SOs was specific to auditory regularities and not to cardiac input.

**Fig. 5.**
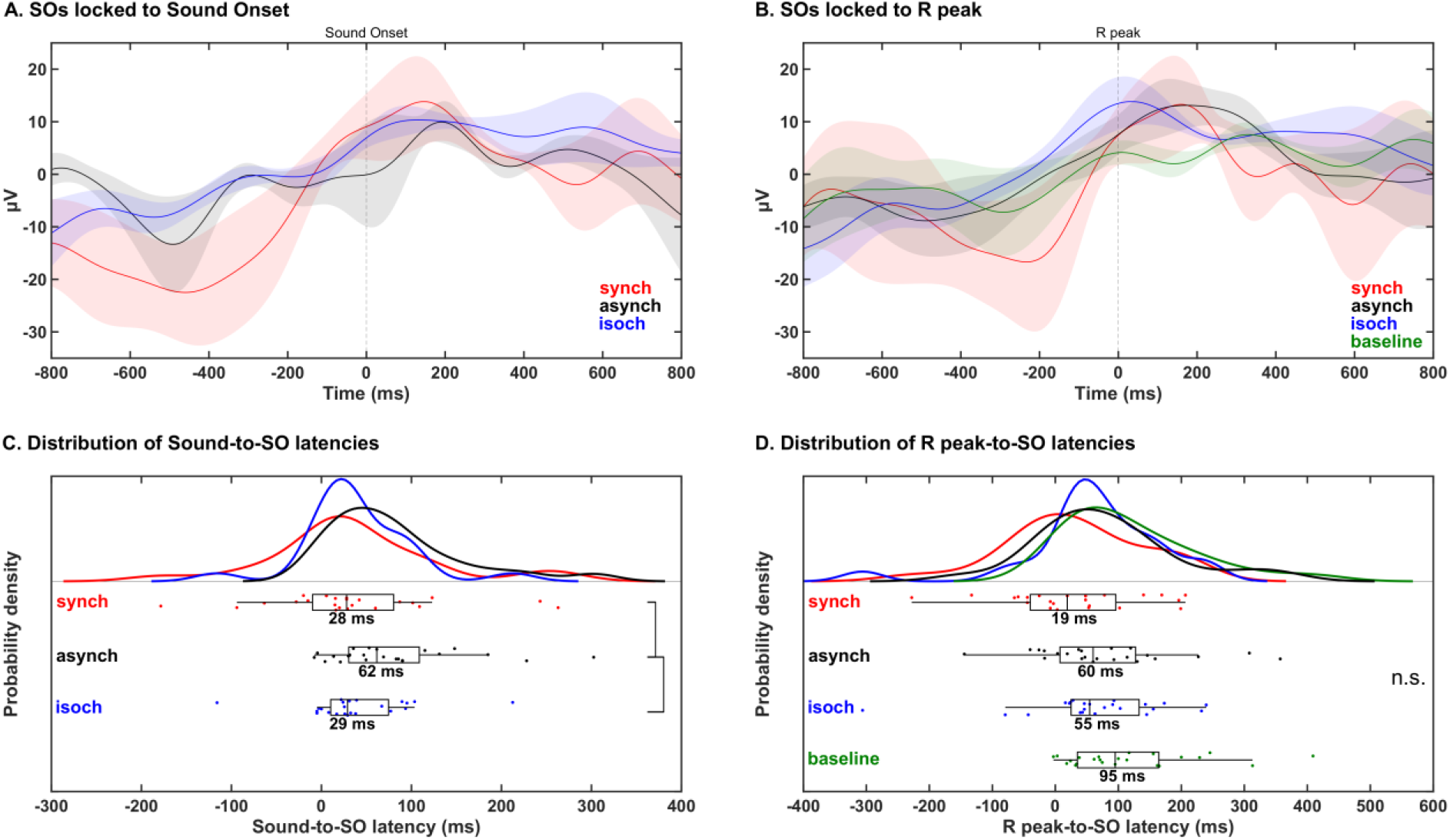
Modulation of SO activity by auditory stimulation in N2 sleep. Grand averaged (N=23) SO waveforms at electrode Cz centred around sound onset (A) and R peak (B) in the synch (red lines), asynch (black lines), isoch (blue lines), and baseline (green lines) conditions during N2 sleep. Grand average and subject-wise sound-to-SO median latency (C) and R peak-to-SO median latency (D) distributions with statistically significant differences (p<0.05; black vertical lines) across conditions. SO peaks are more likely to occur closer to the sound onset in the synch and isoch conditions than in the asynch conditions (C). These results are not trivially driven by a temporal proximity between R peaks and SO peaks in the synch condition (D). n.s. = not significant.

### 5. Quality control analyses

We conducted a series of control analyses on the relation between sound onsets, heartbeat and sound onsets and the heart rate across auditory conditions in order to identify possible factors influencing the neural and cardiac response to sound omissions. Here, unless otherwise specified, we report the results in the cohort of participants included in the ECG analysis, the results of the EEG cohort for wakefulness and N2 sleep being very similar.

#### 5.1. Experimental paradigm control analyses

In the synch condition, the average RS interval was 52.3 ms (SEM = 0.1 ms) for sound trials in wakefulness and sleep, and –2.6 ms (SEM = 2.6 ms) for the isoch and asynch conditions (Fig. S2A). We expected lower RS interval variability in the synch relative to the two other conditions (isoch and asynch), as was confirmed by one way repeated measures Friedman tests with factor Auditory Condition (synch, asynch, isoch) (Fig. S2B; AWAKE: χ^2^ =39.1, p=3.3×10^-9^; N1: χ^2^ =22.5, p=1.3×10^-5^; N2: χ^2^ =36.0, p=1.5×10^-8^; N3: χ^2^ =27.0, p=1.4×10^-6^; REM: χ^2^ =22.5, p=1.3×10^-5^; all post-hoc paired Wilcoxon signed-rank tests showed lower variability in the synch compared to the asynch or isoch condition with p<0.0005). In addition, as expected, the mean SR interval within omission was more variable in the asynch and isoch conditions compared to the synch condition (Fig. S2D; AWAKE: χ^2^ =39.3, p=2.9×10^-9^; N1: χ^2^ =22.5, p=1.3×10^-5^; N2: χ^2^ =36.3, p=1.3×10^-8^; N3: χ^2^ =27.1, p=1.3×10^-6^; REM: χ^2^ =22.8, p=1.1×10^-5^; post-hoc Wilcoxon signed-ranked tests confirmed significant differences with p<0.0005). This first series of control analyses demonstrated that the experimental manipulation, inducing an online fixed temporal alignment between R peak and sound in the synch and a variable one in isoch and asynch, was successful.

The analysis of auditory regularity based on the SS interval confirmed higher variability during the synch and asynch compared to the isoch condition (Fig. S2F; AWAKE: χ^2^_(1,3)_=39.3, p=2.9×10^-9^; N1: χ^2^_(1,3)_=22.5, p=1.3×10^-5^; N2: χ^2^_(1,3)_=37.3, p=7.8×10^-9^; N3: χ^2^_(1,3)_=19.4, p=6.0×10^-5^; REM: χ^2^_(1,3)_=17.7, p=1.0×10^-4^; corroborated by post-hoc Wilcoxon signed-rank tests with p<0.0005) with an average variability value of 0.9 ms in wakefulness and 3.4 ms across sleep stages for the isoch condition (Fig. S2E). This second analysis confirmed that the isoch condition was characterized by highly regular sound-to-sound intervals in comparison to the other conditions.

#### 5.2. Heartbeat control analyses

The heartbeat did not change across experimental conditions during N2, N3 or REM sleep as shown by 1×4 repeated measures Friedman tests on average RR intervals with factor Condition (synch, asynch, isoch, baseline) (Fig. S2G; p>0.05). Conversely, the same analysis performed in wakefulness and N1 sleep revealed significant differences in RR intervals (Fig. S2G; AWAKE: χ^2^ =17.4, p=6.0×10^-4^; N1: χ^2^ =17.0, p=7.0×10^-4^). Post-hoc Wilcoxon signed-rank tests uncovered that significantly reduced RR intervals were specific to the baseline condition (AWAKE: p=0.002 for synch *vs* baseline, p=0.001 for asynch/isoch *vs* baseline; N1: p=0.013 for synch *vs* baseline, p=0.008 for asynch/isoch *vs* baseline) while no differences were observed across auditory conditions (p>0.05).

Finally, for the EEG cohort, we carried out a control analysis on the ECG waveforms to exclude the potential confound of a different degree of contamination of the ECG artifacts in the EEG between conditions of interest upon interpretating the differential EEG signals locked to the R peak. To this aim, we performed a time-wise ECG waveform comparison of the grand averaged ECG trials time-locked to R peaks during sound omissions (Fig. S3). Statistical analysis based on non-parametric cluster permutation statistics (p<0.05, two tailed) contrasting paired experimental conditions (synch, asynch, isoch, baseline) revealed no significant differences in wakefulness and N2 sleep for any of the comparisons.

Overall, these last series of analyses suggest that the heartbeat and ECG characteristics were well matched across auditory conditions and as a consequence, they should not confound the the cardiac and neural differential responses across regularity types.

## Discussion

We investigated the neural and cardiac correlates of cardio-audio regularity encoding in wakefulness and sleep by administering sounds in synchrony to the ongoing heartbeat (synch), at fixed temporal pace (isoch), and in a control condition without specific temporal regularity (asynch) while maintaining matched average sound-to-sound intervals across conditions. We tested whether auditory regularity encoding would result in a prediction error signal as measured by the cardiac and neural signals upon unexpected omitted sounds. A strong cardiac deceleration after sound omission during the cardio-audio sequence suggested an enhanced prediction error signal upon unexpected omissions relative to the other auditory regularity types in all vigilance states. Similarly, neural responses to sound omissions revealed that auditory regularities induced prediction of upcoming sounds both when sounds occurred at a fixed pace and when temporally synchronized to the ongoing heartbeat during wakefulness and N2 sleep. Analysis of the SOs during N2 sleep revealed a reorganization of the ongoing background brain activity both when sounds occurred in synchrony with the ongoing heartbeat and at a fixed temporal pace.

### Cardio-audio regularities result in heart rate deceleration upon sound omissions

The cardiac deceleration upon omission across vigilance states (Fig. 2) can be interpreted in terms of attention reorientation following an unexpected and potentially dangerous event, a parasympathetically-driven effect often reported in conditioning paradigms (45–47). This response can also be associated to a startle reaction and a physiological freezing response, linked to heart rate deceleration, pupil dilation and skin conductance alterations (48–52). In N2 sleep, this cardiac deceleration could be related to cholinergic system engagement. The cholinergic system is known to modulate arousal, with similar mechanisms observed in rats when awakened from NREM sleep by activation of basal forebrain cholinergic neurons using chemical and optogenetic techniques (53, 54). Of note, we found similar deceleration across vigilance states although acetylcholine levels are reduced in N2 sleep and increased in REM sleep compared to wakefulness (55, 56).

Another plausible and not exclusive explanation for the heart rate deceleration following unexpected omission in the synch condition relates to top-down adjustment of cardiac rhythm in order to account for the unexpected silence. As a sound is predicted following a heartbeat (in the synch), the omission may prolong the generation of the next heartbeat within physiologically plausible boundaries so as to ‘wait’ for a delayed auditory stimulus within the sequence, followed by a rapid readjustment to the original rhythm upon subsequent sound presentations. By construction, unlike the synch condition where the heartbeat deceleration was consistently observed, such deceleration was either reduced or absent in conditions where sound presentations bear no temporal relation to ongoing heartbeats (*i.e.* asynch and isoch). In a previous study investigating the cardiac response to sound omission as a function of heartbeat-to-sound onset delay and of interoceptive *vs* exteroceptive attention, a similar cardiac deceleration was reported only in the condition of external attention (28). While this result seems at odds with our finding of preserved cardiac deceleration across vigilance states (and potentially attentional resources), a straightforward comparison is prevented by our lack of control of the focus of attentional resources during wake, also shown to be preserved in sleep (57, 58).

### Neural omission response in cardio-audio and auditory regularity in wakefulness

The neural response to sound omissions was observed as a negative difference in fronto-central electrodes between between the synch and asynch conditions at 165 ms to 218 ms (Fig. 4A) and the isoch and asynch conditions at 114 ms to 159 ms (Fig. 4C). These latencies are well-matched to classic MMN responses observed in wakefulness between 100-200 ms post-stimulus onset (2, 10), upon consideration of the heartbeat to sound latency of 52 ms in the synch condition. Of relevance, the latencies observed here parallel findings from our previous investigation of the synch *vs* asynch comparison where significant differences were observed in the synch *vs* asynch comparison at overlapping latencies between 153-278 ms after R peak onset (29). Our results also resemble previous omission responses (12) peaking at ∼170 ms and arising from the comparison of omission responses with different level of predictability within auditory sequences. Other reports on the neural correlates of predictive processing in auditory regularities have explored different aspects of the omission response. SanMiguel and colleagues investigated the differential responses to button presses that resulted in sounds or omissions and showed similar N100 responses and cortical sources to unpredictable sounds *vs* unpredictable omissions (15), an observation replicated by other groups (59, 60). Finally, while we used largely similar pre-processing (*i.e.* filters) and experiment implementation (*i.e.* online and offline reference) as in Chennu et al. (12), this was not the case for other studies with which direct comparisons are unwarranted.

In our study, the differences between omissions responses and baseline also included earlier components in fronto central electrodes: for the isoch condition starting at 53 ms and for the synch condition starting at 52 ms. We interpret these early differences in light of previous reports suggesting the formation of a sensory template at the predicted sound onset occurring in the same period as the auditory N100 response (61) and localized in the auditory cortex (15, 16). The fact that these early differences did not occur when comparing omissions between auditory regularities (Fig. 4) is likely due to the type of generated prediction. Specifically, upon omissions during auditory regularities, the expectation of a sound is always violated (‘what’ is expected) and therefore the prediction error signal is likely cancelled out in the comparison. The contrast between omissions instead uncovers the prediction error signal of ‘when’ a sound is expected which is generated during isoch and synch conditions. This line of reasoning on the occurrence of an auditory prediction (prediction of ‘what’) in all auditory sequences is also confirmed by the differential response between the omission responses in the asynch *vs* baseline (Fig. S1D).

### Neural omission response in cardio-audio and auditory regularity in N2 sleep

#### Cardio-audio regularity

During N2 sleep, the omission response in cardio-audio regularities (Fig. 4B; synch *vs* asynch comparison) manifested as a long lasting responses unfolding over two time-periods, the first occuring before any sound could be expected (–99-117 ms) and the second taking place at late latencies (322-500 ms). We interpret these findings as the result of the reorganization of the background activity following sound presentation locked to the ongoing heartbeat. This interpretation is supported by two lines of evidence. First, the comparison of sounds to omissions during cardio-audio regularity yielded no significant differences before and around the sound onset period, suggesting that the early difference was driven by auditory stimulus expectation (common to both trials with sound administration and trials with sound omission). Second, cardio-audio encoding induces a reorganization of the background slow oscillatory activity present in N2 sleep, such that the peaks of SOs, periods of maximal excitability, tend to align to the sound onset, when such stimuli can be predicted (as presented in detail in the next section).

#### Auditory regularity

In N2 sleep, we demonstrated a neural response to violation of auditory regularity upon omission of expected sounds. The neural responses during the isoch *vs* asynch condition revealed a posterior negative polarity difference at approximately 100 ms to 200 ms (Fig. 4D). The latencies of this differential response closely matched our results in wakefulness, and, to the best of our knowledge, provides the first account of omission responses in sleep. Previous reports using deviant sound instead of omissions embedded within regular auditory sequences, demonstrated MMN responses in NREM sleep at similar latencies. Importantly, similar to these previous MMN studies in sleep, our omission responses in N2 sleep are characterized by a positive difference at fronto-central electrodes (*e.g.* (62–64)).

### Sound administration modulates SO activity in N2 sleep

The modulation of the synch omission responses in N2 sleep prompted us to look into the possible influence of auditory stimulation on the background oscillatory activity. Indeed, the latencies and topography unveiled in the synch *vs* asynch N2 comparison (*i.e.* fronto-central positivity at –99 ms to 117 ms and fronto-central negativity at 322 ms to 500 ms) could reflect the up and down states of SOs during N2 sleep; keeping in mind that SO peak to trough latencies typically range between 400 ms to 1000 ms (reflecting 0.5-1.2 Hz oscillations; (41–43). Here, this influence was demonstrated by a significantly reduced median latency between the sound stimulus onset and the peak of SOs for the synch and isoch conditions compared to the asynch condition (Fig. 5C). This indicates that sound presentations induced a modulation of slow oscillatory activity in N2 sleep in our subjects in such a way that, when auditory prediction could be generated (*i.e.* synch and isoch), sound onset was more likely to occur close to the SO peaks.

This evidence is reminiscent of the closed-loop auditory stimulation literature wherein temporary synchronization of sound stimuli to ongoing SOs induced an enhancement of the SO rhythm during NREM sleep (39, 43, 44, 65). In the present paradigm as well as in closed-loop auditory stimulation studies, the temporal proximity between sound onset and the SO positive peak suggests the existence of a preferential time window of stimulus processing which may coincide with the positive phase of the SO cycle when neuronal firing is maximal (42, 66, 67). On this basis, auditory regularity encoding would induce a reorganization of the ongoing SO activity, in order to facilitate the neural processing of expected sounds in a sequence when sound onset can be predicted (see also (68) for consistent findings in associative learning bound to the SO peaks).

Of note, not only sounds have been shown to have an impact on the latency of SOs in NREM sleep but also R peaks tend to occur close to the positive peak of the SO compared to other latencies (32, 33). With this in mind, we additionally investigated the potential impact of heartbeat signals on SOs (Fig. 5B,D). In this study, we revealed no significant differences between R peak onset and SO peak across conditions. However, it should be noted that although not significant, we observed a trend of lower R peak to SO peak latencies during cardio-audio regularity compared to the other auditory conditions, possibly driven by the fixed relationship between heartbeat and sound in the synch condition. Overall, these findings suggest that SO latency modulation was specific to auditory regularities and not driven by a systematic temporal readjustmenet of the SOs by the ongoing heartbeat.

### Limitations and future research directions

The selection of a ∼50 ms delay between the detected R peaks and administered sounds led by construction to the investigation of auditory regularity encoding within the systole period in the synch condition. The observed results might be related to this specific temporal period and not necessarily generalize to other latencies of sound administration after the R peak, particularly during the diastole period (19, 69, 70). Future studies will focus on the cardiac and neural correlates of the cardio-audio coupling along different phases of the cardiac cycles.

Another possible limitation relates to the experimental protocol for sleep data acquisition. While participant selection involved an interview assessing each individual’s sleep quality, we did not evaluate our cohort’s general sleep health in a more systematic way using sleep quality assessment questionnaires or actigraphy monitoring, as recommended in sleep research (71). This, along with the auditory stimulation performed and the absence of an adaptation night sleep, could have resulted in a participant cohort with variable sleep quality (72) and have given rise to the low sleep efficiency (69.8%) compared to typically >80% sleep efficiency in healthy unperturbed sleep (*e.g.* (35, 73)). In future work, we will improve participant sleep assessment and sleep quality by including an adaptation night (71) before the data acquisition nights which we speculate will yield improved EEG data availability and will likely result in neural omission responses in the remaining sleep stages (N1, N3, REM), as was observed in the ECG-based cardiac omission response.

### Key contributions

First, we studied the role of heartbeat signals in auditory regularity processing focusing on sound omissions which enabled the investigation of cardio-audio integration free from bottom-up auditory stimulus contributions, differently to previous studies in sleep employing deviant sounds in MMN investigations (*e.g.* (4–6)). Second, the proposed experimental paradigm enabled a counterbalanced ECG artifact between the synch, asynch and baseline conditions when contrasting the OHEPs time-locked to the R peaks, thus allowing the comparison of the neural response to cardiac signals while controlling for heartbeat related artifacts. Third, by investigating both wakefulness and sleep in the same healthy volunteers, we now demonstrate that the human brain infers on the temporal relationship across cardiac and auditory inputs in making predictions about upcoming auditory events across vigilance states. Fourth, we identified a cardiac deceleration as a result of violation detection occurring in parallel to the neural violation response during sound omission. This omission response during N2 sleep was also accompanied by slow oscillation reorganization, representing a possible mechanism through which the brain aligns periods of high neuronal excitability to the expected sound onset.

The cardio-audio synchronicity created *ad hoc* in the experimental environment might reflect real life readjustment of the heartbeat rhythm in order to optimize the temporal relationship between bodily signals and exteroceptive inputs for optimal sensory encoding across vigilance states. These results complement recent accumulating evidence of cardiac signal based markers for assessing the degree of preserved cognitive functioning across a variety of disorders of consciousness (40, 74).

### Conclusion

Collectively, the present results suggest that the human brain can keep track of temporal regularities between exteroceptive inputs, and across interoceptive and exteroceptive inputs during both wakefulness and sleep. Our findings support theories of an interoceptive predictive coding mechanism (20, 21, 75). To our knowledge, this is the first study to investigate auditory regularity processing using omissions and to offer evidence for a potential role of interoceptive inputs under the predictive coding framework in sleep. The conscious and unconscious brain may implicitly process relationships across interoceptive and exteroceptive inputs, allowing the formation of predictions about upcoming sensory events, in order to optimize the signalling and prediction of potential upcoming dangers.

## Materials and Methods

### Ethics statement

Approval for the study (Project-ID: 2020-02373) was obtained by the local ethics committee (La Commission Cantonale d’Ethique de la Recherche sur l’Etre Humain), in accordance with the Helsinki declaration.

### Sample size estimation

The number of participants selected for this study was based on the results of our previous study in wakefulness (29). Sample size was derived from the electrodes within significant clusters and the latencies at which they were observed when comparing OHEPs in the synch *vs* asynch comparison. After fixing the probability threshold for rejecting the null hypothesis to 0.05 (two-tailed), we simulated 5000 random replications of the neural responses to sound omissions. These simulations were based on extracting random samples from a multivariate normal distribution with mean and variance estimated from previous data (in accordance with https://osf.io/rmqhc/, documented in the publically available Fieldtrip toolbox, https://www.fieldtriptoolbox.org/example/samplesize/). Cluster-based permutation statistical analysis on each of the simulated comparisons, provided a power > 0.90 (i.e. the percentage of times the results were significant) with a sample size higher than seventeen for the healthy cohort (N>17). Here, we decided to recruit twenty-six volunteers, in order to account for possible participant exclusion due to a higher likelihood of equipment malfunction, excessive artifactual trials or channels during full-night sleep recordings.

### Human participants

Twenty-six self-reported good sleeper volunteers took part in both the wakefulness and sleep arms of this study. Participants were considered eligible if they had no history of psychiatric, neurological, respiratory or cardiovascular conditions, no sleep apnoea, and a regular sleep schedule, evaluated during a phone interview. Hearing conditions were an additional exclusion criterion. All participants gave written informed consent and received approximately 150 Swiss Francs as monetary compensation.

### Experimental design

A two-way crossover experimental design was implemented in this study. Participants attended one wakefulness and one sleep session on two occasions separated by a minimum of one day and a maximum of ten days (wakefulness session first for 12 out of 26 participants). In both sessions, participants were instructed to passively listen to the administered sound sequences. They were naïve to the experimental manipulation, as suggested by informal verbal inquiry regarding the experimental design after the experiment. At the end of the second session, and if desired, participants were debriefed about the purpose of the experiment.

The sleep session recordings took place in a sound-attenuated hospital room equipped with a comfortable hospital bed to allow for overnight sleep recordings. During the sleep session, participants arrived at the laboratory at approximately 9 pm and following set-up preparation, they were instructed to lie down and inform the experimenters when they were ready to sleep. Lights were then switched off, the auditory stimulus administration commenced and the volunteers were left alone to naturally fall asleep. Although participants had the liberty to leave at any time, we explained that their inclusion in the study required a minimum of four hours of continuous data acquisition after sleep onset. They were free to choose to spend the night at the sleep laboratory and to be woken up at a desired time or by approximately 7 am the next morning, at which time lights were switched on.

In both sessions, participants were equipped with electrodes for heartbeat (ECG), eye movement (EOG), and EEG recordings (see below). For the sleep session, additional electrodes for submental electromyography (EMG) were attached, in accordance with the 2007 AASM guidelines for sleep scoring (76). In-ear phones (Shure SE 215, Niles, IL) were utilized during both sessions instead of external headphones, in order to increase sound attenuation, subject comfort during sleep and to prevent physical contact with and thus displacement of EEG cap and electrodes. The online EEG, ECG, EOG and EMG were continuously monitored by the experimenters to ensure effective stimulus administration, data acquisition, heartbeat detection and sleep quality throughout both sessions.

### Stimuli

Sound stimuli were 1000 Hz sinusoidal tones of 100 ms duration (including 7 ms rise and fall times) and 0 μs inter-aural time difference. A 10 ms linear amplitude envelope was applied at stimulus onset and offset to avoid clicks. Stimuli were 16-bit stereo sounds sampled at 44.1 kHz and were presented binaurally with individually adjusted intensity to a comfortable level for wakefulness. A considerably lower than wakefulness intensity of approximately 45 dB was chosen for sleep, in order to facilitate a non-fragmented sleep session without multiple awakenings.

### Experimental procedure

During the wakefulness session, volunteers sat comfortably on a chair in a sound-attenuated experimental room and were instructed to keep their eyes open, avoid excessive eye blinking, body and jaw movements; the aforementioned measures served in ensuring high signal quality. Each participant was presented with four types of stimulation conditions administered in separate experimental blocks in a pseudo-random order and was asked to passively listen to the sounds while keeping the eyes fixed on a cross centrally located in the visual field. The conditions were a baseline without auditory stimulation and three auditory conditions, namely synch, asynch and isoch. During wakefulness, the baseline lasted ten minutes and was acquired prior to auditory stimulation. During sleep, numerous two-minute baseline blocks were acquired in alternation to the sound sequences in an attempt to ensure that baseline background activity was comparable to the preceding sound stimulation during all stages of sleep. The three auditory conditions lasted five minutes each and corresponded to separate experimental blocks, which were repeated six times during wakefulness in a semi-randomized order. During the sleep session, sounds were administered for the entire length of the sleep recording in sequences of three auditory blocks always followed by a baseline (*e.g.* isoch-synch-asynch-baseline or synch-asynch-isoch-baseline).

### Auditory conditions

All auditory conditions consisted of the sequential presentation of 250 stimuli (80% sounds and 20% omissions) administered in a pseudo-random order wherein at least one sound stimulus intervened between two subsequent omissions. Details for each auditory condition are given below and a thorough post-hoc evaluation of the experimental manipulation is provided in the Results and Supporting Information (Results 5. Quality control analyses & Fig. S2).

In the synch condition, the temporal onset of each sound stimulus was triggered by the online detection of R peaks from raw ECG recordings. To enable effective online R peak detection, raw ECG recordings were analyzed in real-time using a custom MATLAB Simulink script (R2019b, The MathWorks, Natick, MA). The variance over the preceding 50 ms time window was computed and an R peak was detected when the online ECG value exceeded an individually adjusted 10–15 mV^2^ variance threshold, which in turn triggered the presentation of a sound stimulus or an omission. This procedure resulted in a fixed R peak-to-sound average delay (*i.e.* RS interval) of 52 ms (SD = 5 ms) for wakefulness and sleep across participants, the minimum fixed delay offered by the utilized equipment in order to best approximate the condition of co-occurrence between the heartbeat and auditory stimuli over time.

In the asynch condition, the onset of sound presentation was based on the RR intervals extracted from a previously acquired synch block. Specifically, the ECG recorded during the preceding synch block was analyzed offline to extract RR intervals by automatic detection of R peaks and computation of RR intervals. 250 RR intervals were selected if they were above the 25^th^ and below the 75^th^ percentile of RR interval distribution in the synch block, in order to take into account possible missed R peaks in the online detection during the synch block. Next, RR interval order was shuffled giving rise to a predefined pseudo-random sequence closely resembling the participant’s heartbeat rhythm. By construction, differences between the synch and asynch conditions in terms of average and variance of the RR intervals were minimized, contrary to the RS interval being fixed in the synch condition and variable in the asynch condition.

In the isoch condition, the onset of sound presentations was based on the median RR interval calculated during a previously acquired synch block. This procedure produced similar sound-to-sound intervals across the synch, asynch and isoch conditions however, unlike the synch and asynch conditions, sound-to-sound intervals in the isoch condition had low variability.

### Data acquisition

Continuous EEG (g.HIamp, g.tec medical engineering, Graz, Austria) was acquired at 1200 Hz from 63 active ring electrodes (g.LADYbird, g.tec medical engineering) arranged according to the international 10–10 system and referenced online to the right earlobe and offline to the left and right ear lobes. Electrode AFz served as the ground. Biophysical data were acquired using single-use Ag/AgCl electrodes. Three-lead ECG was recorded by attaching two electrodes (a third was a reference) to the participant’s chest on the infraclavicular fossae and below the heart. A vertical EOG electrode was attached below the right eye and a horizontal EOG electrode was attached to the outer right canthus. Since muscle atonia is associated with increased sleep depth and is an essential marker for effective sleep staging (76), EMG was additionally acquired sub-mentally during the sleep session alone. Impedances of all active electrodes were kept below 50 kΩ. All electrophysiological data were acquired with an online band-pass filter between 0.1 and 140 Hz and a band-stop filter between 48 and 52 Hz to reduce electrical line noise.

### Sleep scoring

An experienced sleep scoring specialist (Somnox SRL, Belgium), blind to the experimental manipulation in this study, performed the scoring of the continuous sleep electrophysiological data in order to pinpoint the periods of wakefulness and micro-arousal as well as periods of N1, N2, N3 and REM sleep. Sleep scoring was performed via visual inspection of contiguous 30-second segments of the EEG, EOG and EMG time-series, as outlined in the 2007 AASM guidelines for sleep scoring (76). Segments scored as periods of wakefulness or micro-arousals in the sleep recordings were excluded from further analysis.

### Data analysis

Electrophysiological data analyses were performed in MATLAB (R2019b, The MathWorks, Natick, MA) using open-source toolboxes EEGLAB (version 13.4.4b, (77)), Fieldtrip (version 20201205, (78)), as well as using custom-made scripts. Raincloud plots were generated using the Raincloud plot toolbox (79).

### R peak detection

The R peaks in the continuous raw ECG signal were selected offline using a semi-automated approach as in Pfeiffer & De Lucia (29). The custom-made MATLAB script, *rpeakdetect.m* (https://ch.mathworks.com/matlabcentral/fileexchange/72-peakdetect-m/content/peakdetect.m), was utilized to automatically identify the sharp R peaks in the raw ECG signal. Visual inspection of the online and offline detected R peaks ensured that the selected peaks fitted within the expected structure of the QRS complex in the continuous raw ECG signal. Frequent flawed online identification of the R peaks or faulty auditory stimulus presentation in a given block resulted in the exclusion of the block from a given participant’s dataset. For blocks that were included, unrealistic RR, RS and SR interval values, observed as a result of infrequent flawed offline marking of R peaks, were identified and excluded using the *rmoutliers* MATLAB function (R2019b, The MathWorks, Natick, MA) and visual inspection of the detected R peaks and selected outliers.

### Quality control analyses

In the following, we will refer to R as the latency of the R peak of the ECG waveform, S represents the sound onset latency, RS the R peak-to-sound latency, SR the sound to the next R peak latency, SS the sound-to-sound latency, RR the R peak-to-R peak latency and interval variability is calculated as the standard error of the mean (SEM). A series of control analyses were performed to investigate whether the experimental manipulation was producing the expected RS, SR, RR and SS mean and variances, and the presence of possible confounding factors. SS and RR intervals (for sound trials preceded by sound trials) were extracted for the synch, asynch, isoch and baseline conditions where relevant. In addition, RS intervals for sound trials and SR intervals for omission trials were computed to quantify the degree of cardio-audio synchronization and heartbeat onset variability during sound omission, respectively. Variability in the same interval measures, computed as the SEM was additionally investigated. Of note, unlike RR, SR and RS variability, SS variability was first computed within a given experimental block and then across experimental blocks, to account for ongoing changes in RR intervals (and hence SS intervals) in sleep. Non-parametric one way repeated measures Friedman tests (p<0.05) were performed on the average RS, SR, SS and RR intervals and variabilities of each participant for wakefulness and all sleep stages with within-subject factor Condition (3 levels for RS, SR, SS intervals and variability: synch, asynch, isoch; 4 levels for RR intervals and variability: synch, asynch, isoch, baseline). Post-hoc paired Wilcoxon signed-rank tests (p<0.05) identified any significant pairwise comparisons (no multiple comparisons correction was applied since pairwise differences were of interest).

To ensure no significant differences in the ECG signal across experimental conditions, ECG waveforms during omissions were extracted from ECG recordings between –100 ms and 500 ms relative to R peak onset, matching the EEG trial-based analysis (see below, EEG Data Analysis). The non-parametric cluster-based permutation statistical analysis approach (80) was employed to investigate ECG waveform differences between the various experimental conditions outlined herein. In order to reject the null hypothesis that no significant differences existed in the given set of experimental conditions being contrasted, maximum cluster-level statistics were determined by shuffling condition labels (5000 permutations), allowing for a chance-based distribution of maximal cluster-level statistics to be estimated. Since maxima are utilized by this method, it enables the correction of multiple comparisons over time. A two-tailed Monte-Carlo p-value allowed for the definition of a threshold of significance from the distribution (p<0.05, two-tailed).

### ECG data analysis

The identified R peaks were used to derive omission trial related RR intervals. The representative R peak for an omission trial was selected as the first R peak following an omission in the continuous ECG signal. The omission RR Interval (RR_om_) was therefore calculated as the latency between the selected omission R peak and the R peak immediately preceding it. In order to investigate potential heartbeat alterations associated with the omission trial, additional RR intervals were identified using contiguous R peaks to reflect the RR interval for the trial prior to omission (RR_-1_) and up to two trials following omission (RR_+1_ and RR_+2_). Omission trials were considered only if they were followed by at least two sound stimuli to ensure no overlap between investigated RR intervals.

The 30-second sleep stage labeled epochs were used to label omission trials for all auditory conditions. For the analysis of the ECG signals during the wakefulness and sleep sessions, we imposed a minimum of 30 artifact-free trials for the ECG data analysis and for each condition of interest. Hence, different sets of RR intervals formed the final dataset per participant and consisted of five vigilance states: AWAKE, N1, N2, N3 and REM sleep where available (Table S1-S2). One way ANOVAs on RR interval quantities showed no significant differences (p>0.05) in trial numbers across auditory conditions within each vigilance state, therefore all RR intervals were included in the statistical analysis. RR intervals for each omission trial order type were averaged for each auditory condition, participant, and each vigilance state.

### ECG statistical analysis

First, 3×4 repeated measures ANOVAs were performed on the average RR intervals with within-subject factors Auditory Condition (synch, asynch, isoch) and Trial Order (one trial before, trial during, first trial after, second trial after sound omissions) for the wakefulness session (AWAKE) and each of the sleep stages (N1, N2, N3, REM) separately. Second, for a sub-set of participants (N=6) who had sufficient trials across wakefulness and all sleep stages, a 5×3×4 repeated measures ANOVA was performed on the average RR intervals with within-subject factors Vigilance State (AWAKE, N1, N2, N3, REM), Auditory Condition (synch, asynch, isoch) and Trial Order (one trial before, trial during, first trial after, second trial after sound omission). Of note, unlike comparisons within vigilance states, trial numbers were matched across vigilance states by randomly selecting a subset of RR intervals based on the minimum number (≥30 trials) of available RR intervals across participants to ensure accurate comparison across vigilance states. In order to reject the null hypothesis that no significant differences existed in the given set of ANOVA factors being compared, each of the factors’ condition labels were shuffled (2000 permutations), allowing for an estimation of the chance-level distribution of maximal cluster-level statistics to be estimated, based on non-parametric statistical testing. Wilcoxon signed-ranked tests (p<0.05) were used to evaluate whether significant differences existed between the distribution of post-permutation F values and the true main effect and interaction effect F values. Post-hoc Wilcoxon signed-rank tests with Bonferroni correction for multiple comparisons (p<0.018 for auditory condition and p<0.013 for omission trial order comparisons) were utilized for pairwise comparisons between all investigated within-subject variables.

### EEG data analysis

Continuous raw EEG data were band-pass filtered using second-order Butterworth filters between 0.5 and 30 Hz for the wakefulness and sleep session. To look specifically at slow wave activity, raw electrophysiological data were band-pass filtered between 0.5 and 4 Hz, a well-established frequency range for investigations of slow-wave activity in the delta band range (81). Independent Component Analysis (ICA) as implemented in Fieldtrip (77, 78) was used in order to identify and remove any eye movement activity related components in the continuous wakefulness and sleep EEG.

Wakefulness and sleep continuous EEG data were epoched between –100 ms and 500 ms relative to the selected event onset. This range was chosen in reflection of previously reported classic omission response latencies ((12) and others) and importantly, to ensure no overlap between consecutive auditory stimulation trials (as in (29), assuming a range of 60-100 beats per minute across volunteers). Various event onsets were selected, namely, heartbeat onset, sound onset and omission onset, depending on the comparison performed (Results 3. Neural omission response). Of note, while the isoch condition had fixed SS intervals, this was not the case in the asynch condition. To test whether sound prediction was based on computing the time of maximal sound occurrence probability, average sound-based omission responses were calculated by extracting epochs from the continuous EEG asynch and isoch recordings, now time-locked to the latency occurring during the omission at the average SS interval and resulting in the estimation of the OEP.

The resting-state baseline recordings acquired as part of the experimental procedure (without auditory stimulation) were utilized to extract EEG evoked potentials with matched event onsets to auditory stimulation trials and epoched between –100 ms and 500 ms. First, we extracted HEPs based on heartbeat onset in the resting state for upcoming statistical comparisons to the synch and asynch OHEP. Second, in the absence of sound stimulation, the baseline allowed for comparing the OEPs of the isoch and asynch conditions to a control condition. In this case, a random selection of epochs was extracted from continuous baseline recordings, such that the latencies between epoch onset and closest heartbeat (*i.e.* R peak) were matched on the single-trial level to the trial onsets in the sound-based isoch and asynch conditions. Upcoming pre-processing steps were replicated for auditory stimulation and baseline trials, for each participant, and for wakefulness and all sleep stages as follows.

Artifact electrodes and trials were identified using a semi-automated approach for artifact rejection as implemented in Fieldtrip (78). Noisy EEG electrodes were excluded based on a signal variance criterion (3 z-score Hurst exponent) and substituted with data interpolated from nearby channels using spherical splines (82). Across eligible participants, an average of 8.0 (SD = 2.3) electrodes (M = 12.9%, SD = 3.7%) were interpolated in wakefulness and an average of 8.9 (SD = 1.1) electrodes (M = 14.4% SD = 1.7%) were interpolated in sleep. In wakefulness data, trials containing physiological artifacts (*e.g.* eye movement, excessive muscle activity), not accounted for by ICA, were identified by visual inspection and by using a 70 μV absolute value cut-off applied to the EEG signal amplitude and were excluded from further analysis. A higher absolute value cut-off of 300 μV was employed for the selection of artifactual epochs in sleep, in order to prevent the exclusion of high-amplitude slow wave activity (83). Extreme outliers in signal kurtosis and variance were additional criteria for the rejection of artifactual trials from sleep recordings. Finally, common average re-referencing was applied.

The 30-second sleep stage labeled epochs were used to label artifact-free trials for all event onset types and for all three auditory conditions and the baseline. Hence, five different sets of trials formed the final processed dataset per participant for each of the five vigilance states: AWAKE, N1, N2, N3 and REM sleep where available (Table S1-S2). We note that we chose not to employ pre-stimulus baseline correction in wakefulness or sleep trials. In addition, since one way ANOVAs on artifact-free trial numbers for each evoked response of interest confirmed that no significant differences (p>0.05) existed across auditory conditions alone, all trials were kept in auditory condition comparisons. Conversely, since acquired baseline data were significantly less (p<0.05) than the three auditory conditions, baseline trial numbers were quantitatively matched for each participant and for each vigilance state between auditory conditions and baseline. The number of available trials for each vigilance state, averaged across experimental conditions and event onsets of interest were AWAKE: M = 286, SD = 13 trials; N1: M = 97, SD = 22 trials; N2: M = 428, SD = 184 trials; N3: M = 159, SD = 72 trials; REM: M = 163, SD = 65 trials. Finally, grand average evoked responses to sound omissions were derived.

### EEG statistical analysis

EEG statistical analyses were performed in MATLAB (R2019b, The MathWorks, Natick, MA) using Fieldtrip (version 20201205, (78)), as well as custom made scripts. For the analysis of the electrophysiological signals during the wakefulness and sleep sessions, we imposed a minimum of 60 artifact-free trials for the EEG data analysis, chosen based on the signal-to-noise ratio required to meaningfully interpret EEG statistical analysis results (84).

The non-parametric cluster-based permutation statistical analysis approach (80) was employed to investigate sensor-level EEG-based differences between the various experimental conditions outlined herein. Under this statistical framework, statistically significant individual data samples were grouped based on the degree of their shared spatial and temporal characteristics. The resulting clusters were statistically evaluated by summating the t-values for all samples forming up a given cluster. In order to reject the null hypothesis that no significant differences existed in the given set of experimental conditions being contrasted, maximum cluster-level statistics were determined by shuffling condition labels (5000 permutations), allowing for a chance-based distribution of maximal cluster-level statistics to be estimated. A two-tailed Monte-Carlo p-value allowed for the definition of a threshold of significance from the distribution (p<0.05, two-tailed). Of note, the cluster permutation based multiple comparisons correction only applied across channels and latencies when comparing two experimental conditions, however no multiple comparisons correction was applied across the number of comparisons made in this study. In wakefulness and sleep, this procedure was performed over the entire trial length from –100 ms to 500 ms relative to the event onset of interest (heartbeat onset for OHEP comparisons and expected sound onset based on average SS interval for OEP comparisons). Finally, in order to evaluate the size of the observed effects, the Cohen’s d statistic was calculated at the peak latenies of significant clusters (85).

### SO data analysis

To examine whether N2 sleep EEG statistical analysis results could have been influenced by the relationship between sounds and SO activity (39, 43, 65) and/or between heartbeats and SO activity (32) in NREM sleep, we identified SOs in N2 sleep artifact-free EEG data. We then computed the latency at which the SOs occurred compared first, to sound presentations in the auditory conditions and second, to heartbeats in the auditory conditions and baseline.

SOs in the continuous EEG time-series were detected across experimental conditions over frontocentral electrode Cz where a high probability for SO detection is to be expected (41). We marked SOs based on the method described in Ngo *et al.* (43) and in Besedovsky *et al.* (65).

In brief, the 0.5 to 30 Hz band-pass filtered data were downsampled from 1200 Hz to 100 Hz and a low pass finite impulse response filter of 3.5 Hz was used in order to improve the detection of SO components. Next, N2 sleep labeled segments were extracted for the identification of SOs. Consecutive positive-to-negative zero crossings were picked out and were selected only if their temporal distance was between 0.833 s and 2 s, yielding a frequency between 0.5 Hz and 1.2 Hz for the designated oscillations. Negative and positive peak potentials in the oscillations were defined as the minima and maxima present within eligible consecutive positive-to-negative zero crossings. The mean negative and mean positive-to-negative amplitude differences were calculated across selected oscillations at electrode Cz. For each identified oscillation that satisfied the frequency criterion, if the negative amplitude was 1.25 times lower than the mean negative amplitude and the positive to negative amplitude difference was 1.25 times higher than the mean positive to negative amplitude difference, the oscillation was marked as a SO.

The positive half-wave peak time point was chosen as the representative latency for each SO in light of relevant literature (42, 66) and following visual inspection of the SO and sound presentation time-series. Sound to SO latency for the synch, asynch and isoch conditions and R peak to SO latency for the synch, asynch, isoch and baseline conditions were computed for all artifact-free sound trials. Omission trials were excluded from this analysis since in this case, we were interested in how sounds and the sound to heartbeat relationship may modulate SOs in N2 sleep. Latencies were considered as valid only if they were between –800 and 800 ms (32), a range chosen to minimize potential contamination of the sound or R peak to SO relationship by upcoming sounds or heartbeats. Median latencies were calculated for each subject and condition separately.

### SO statistical analysis

In N2 sleep, a non-parametric 1×3 repeated measures Friedman test was calculated on the median sound to SO latencies at electrode Cz with within-subject factor Auditory Condition (synch, asynch, isoch) and a 1×4 repeated measures Friedman test were computed on the median heartbeat to SO latencies with within-subject factor Condition (synch, asynch, isoch, baseline). Pairwise comparisons were calculated using post-hoc paired Wilcoxon signed-rank tests between all investigated within-subject variables (no multiple comparisons correction was applied since pairwise differences were of interest). Latency trial numbers were matched for the median heartbeat to SO latency comparison but not for the sound to SO latency comparison, since in the former, baseline recordings lengths were significantly lower to auditory condition recordings (as confirmed by 1×4 and 1×3 repeated measures ANOVAs on latency trial numbers, which significantly differed only across heartbeat to SO trials and not sound to SO trials).

### Data availability

All the data used to generate the main figures are available for review at https://osf.io/hmqza/?view_only=45a3a0c8a73c4cafa7f5d3201777d813 and will be made publically available upon publication. The data that were not used in the main figures will be shared upon request.

### Code availability

Costum-made code used to run the quality control analyses and EEG and ECG data analyses are available on https://github.com/DNC-EEG-platform/CardioAudio_Sleep/. Any additional information required to reanalyze the data reported in this paper is available from the corresponding authors upon request.

## Acknowledgments & Funding Sources

We express our gratitude to Nathalie Nguepnjo Nguissi, Elsa Dosi and Diana Ortolani for assistance during data acquisition. We extend our thanks to Dr Aurore Guyon Postalci from SOMNOX for performing the stage scoring of the sleep data. We thank Rupert Ortner and the g.tec team for technical support.

This work was supported by a Spark SNSF Grant (no. 196194) and the Catalyst Fund of Bertarelli Foundation awarded to M.D.L. Additional funding was provided by an SNSF Grant (no. 320030_182589) awarded to S.S.

