## supplementary_material for "Cardio-audio synchronization elicits neural and cardiac surprise responses in human wakefulness and sleep"

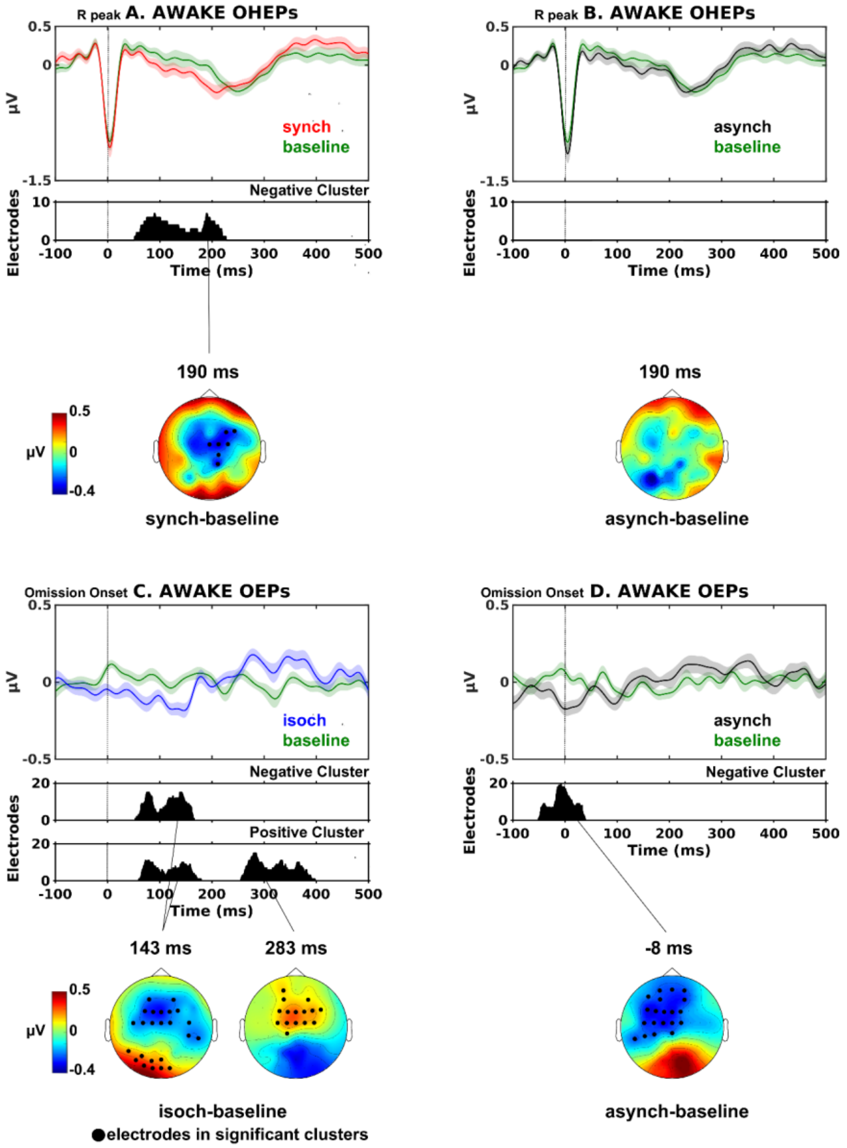


**Fig. S1.** Neural omission response in wakefulness. OHEPs and OEPs comparisons during sound omissions in auditory conditions contrasted to the baseline (green lines) during wakefulness (AWAKE; N=23). The synch vs baseline comparison (A) yielded a significant negative cluster ($p$ = 0.018, Cohen's d = 1.026) on central scalp electrodes at 52 ms to 227 ms. The asynch vs baseline comparison (B) showed no significant differences. (A) & (B) suggest that only in the synch condition, the cardio-audio regularity induced a modulation of OHEPs with respect to baseline in wakefulness. The isoch vs baseline comparison (C) revealed a significant negative cluster ($p = 0.008$, Cohen's $d = 0.783$) at 53 ms to 166 ms following expected sound onset on anterior-central scalp electrodes and a significant positive cluster ($p = 0.030$, Cohen's $d = 0.641$) at similar latencies of 60 ms to 179 ms in posterior electrodes. An additional positive cluster ($p = 0.005$, Cohen's $d = 1.214$) was identified in the same comparison at 256 ms to 398 ms on anterior-central electrodes. The asynch vs baseline comparison (D) also resulted in a significant negative cluster ($p = 0.004$, Cohen's $d = 0.923$) on anterior-central scalp electrodes at -51 ms to 38 ms following expected sound onset. (C) & (D) provide evidence of an auditory violation response both in the regular isoch and pseudo-regular asynch conditions.

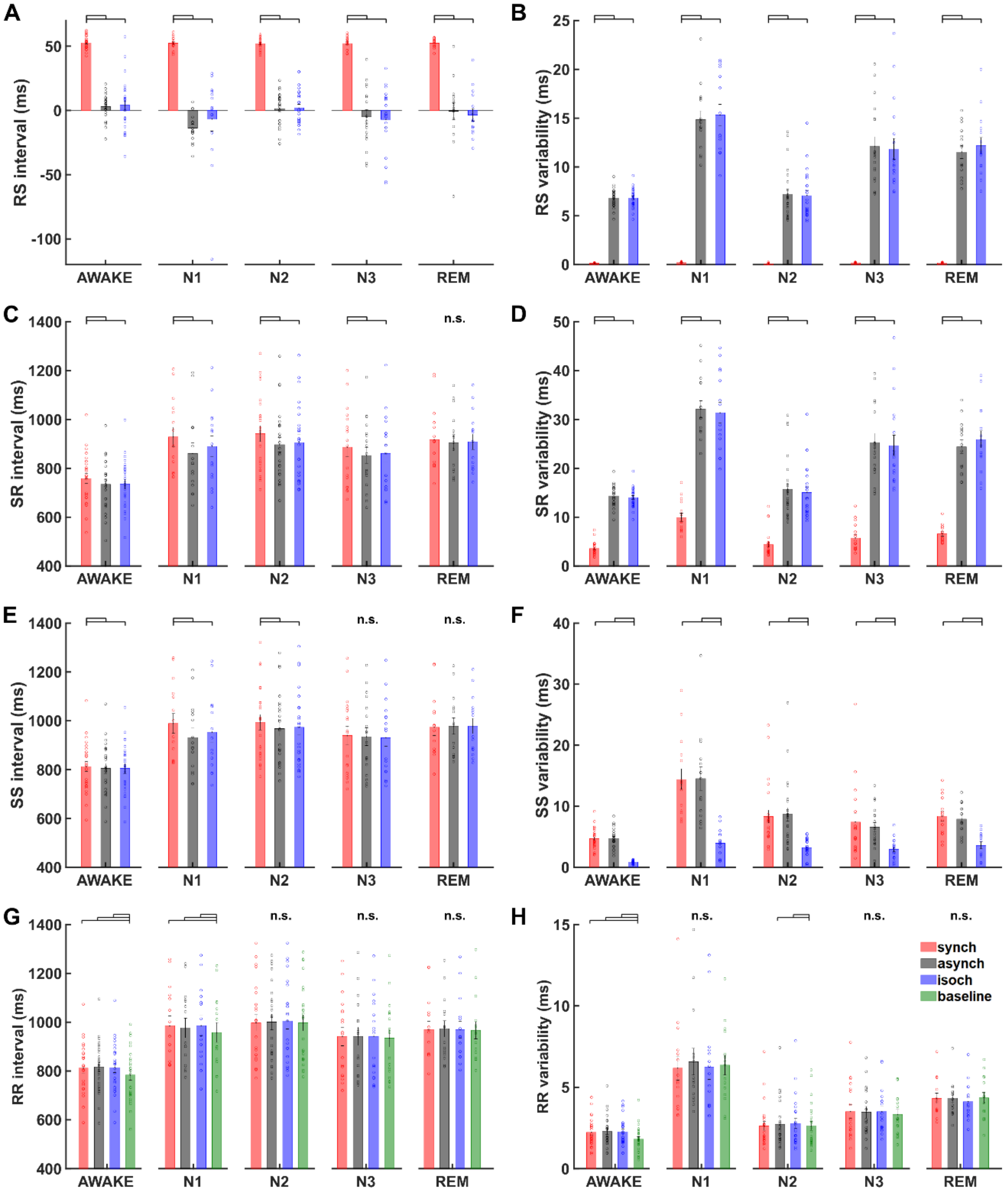


**Fig. S2.** Quality control analyses across wakefulness and sleep. Single-subject and grand averaged RS interval (A) and variability (B), SR interval (C) and variability (D), SS interval (E) and variability (F), RR interval (G) and variability (H) values and statistical analysis results for all auditory regularity conditions (synch: red; asynch: black; isoch: blue; baseline: green) and for wakefulness (AWAKE; N=26) and all stages of sleep (N1, N=15; N2, N=24; N3, N=18; REM, N=15). Variability refers to the SEM of corresponding intervals. As expected, RS and SR variability were significantly lower in the synch condition compared to the asynch and isoch conditions. SS variability was significantly lower in the isoch condition compared to the synch and asynch conditions. RR intervals were not significantly different across experimental conditions in N2, N3 and REM sleep and not significantly different across auditory conditions in wakefulness and N1 sleep. Black horizontal lines indicate statistically significant ($p<0.05$) differences across experimental conditions. Error bars indicate the ±SEM.

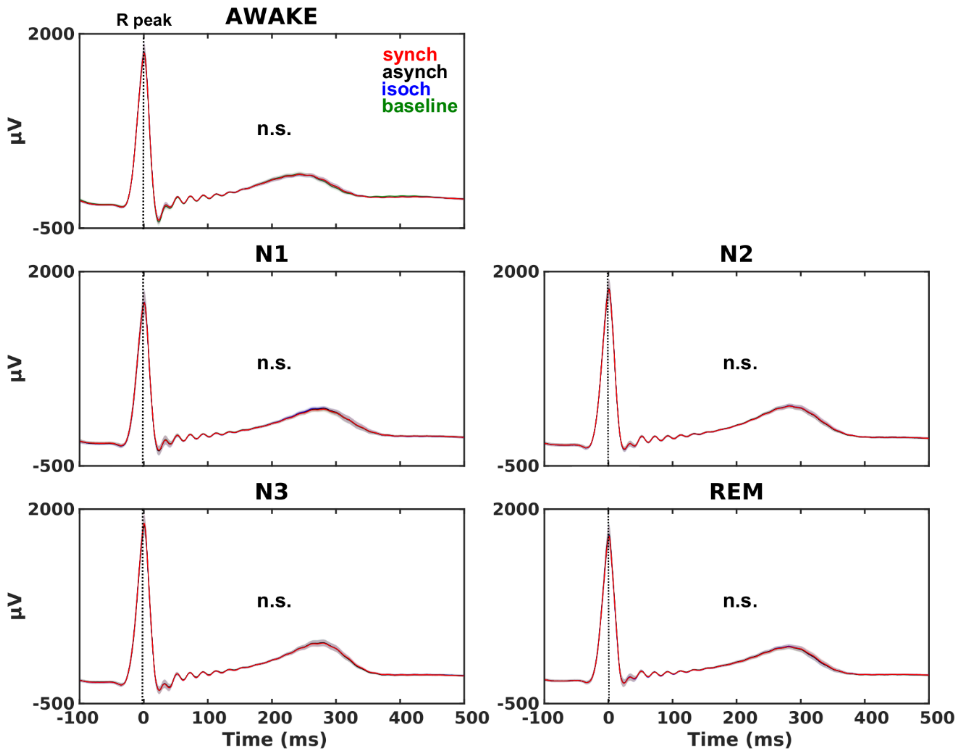


**Fig. S3.** ECG waveforms in wakefulness and sleep. Grand averaged ECG waveforms (time-locked to the closest R peak during omissions) in the synch (red lines), asynch (black lines), isoch (blue lines) and baseline (green lines) experimental conditions for the wakefulness session and all stages of sleep. Shaded regions indicate ±SEM across participants. Cluster permutation statistical analysis ($p<0.05$, two tailed) contrasting experimental conditions revealed no significant differences in the ECG signal during omissions in wakefulness and any sleep stage.

**Table S1.** Participants included in each sleep stage and for each type of analysis. Total participant numbers in the cardiac omission response analysis (ECG) and neural omission response analysis (EEG) for wakefulness (AWAKE), sleep (ALL) and each of the stages of the sleep cycle (N1, N2, N3, REM). Participant exclusion was based on a threshold of available trials to allow for sufficient signal to noise ratio in the EEG analysis (≤60 trials) and for sufficient statistical power in the ECG analysis (≤30 trials).

| | **AWAKE** | **SLEEP** | | | | |
|---|---|---|---|---|---|---|
| Participants | | **ALL** | **N1** | **N2** | **N3** | **REM** |
| **ECG** | 26 | 25 | 15 | 24 | 18 | 15 |
| **EEG** | 23 | 25 | 12 | 23 | 14 | 13 |

**Table S2.** List of participants included in each vigilance stage. Participant-specific inclusion in the cardiac omission response statistical analysis (ECG) and in the neural omission response statistical analysis (EEG) for wakefulness (AWAKE) and each of the stages of the sleep cycle (N1, N2, N3, REM, where relevant). √ denotes participant inclusion in the analysis and × denotes participant exclusion from the analysis.

| | **ECG** | | | | | **EEG** | |
|---|---|---|---|---|---|---|---|
| | AWAKE | N1 | N2 | N3 | REM | AWAKE | N2 |
| **s1** | √ | × | × | √ | × | x | × |
| **s2** | √ | × | × | × | × | √ | × |
| **s3** | √ | √ | √ | × | √ | √ | √ |
| **s4** | √ | √ | √ | × | √ | √ | √ |
| **s5** | √ | √ | √ | √ | √ | √ | √ |
| **s6** | √ | × | √ | × | √ | √ | √ |
| **s7** | √ | √ | √ | × | × | √ | × |
| **s8** | √ | √ | √ | √ | × | √ | √ |
| **s9** | √ | √ | √ | √ | √ | x | √ |
| **s10** | √ | × | √ | × | × | √ | √ |
| **s11** | √ | √ | √ | √ | √ | √ | √ |
| **s12** | √ | × | √ | √ | × | √ | √ |
| **s13** | √ | √ | √ | √ | √ | √ | √ |
| **s14** | √ | √ | √ | √ | √ | √ | √ |
| **s15** | √ | √ | √ | √ | × | √ | √ |
| **s16** | √ | × | √ | √ | √ | √ | √ |
| **s17** | √ | √ | √ | √ | × | √ | √ |
| **s18** | √ | √ | √ | × | × | √ | √ |
| **s19** | √ | × | √ | √ | √ | √ | √ |
| **s20** | √ | √ | √ | × | √ | √ | √ |
| **s21** | √ | × | √ | √ | √ | √ | √ |
| **s22** | √ | √ | √ | √ | √ | √ | √ |
| **s23** | √ | √ | √ | √ | × | √ | √ |
| **s24** | √ | × | √ | √ | × | √ | √ |
| **s25** | √ | × | √ | √ | √ | √ | √ |
| **s26** | √ | × | √ | √ | √ | × | √ |
